# Small-scale commercial chicken production: A risky business for farmers in the Mekong Delta of Vietnam

**DOI:** 10.1101/2020.11.02.364281

**Authors:** Truong Dinh Bao, Nguyen Van Cuong, Hoang Doan Phu, Nguyen Thi Thuy Dung, Bach Tuan Kiet, Jonathan Rushton, Juan Carrique-Mas

## Abstract

Small-scale farming of meat chicken flocks using local native breeds contributes to the economy of many rural livelihoods in Vietnam and many other low- and middle-income countries (LMICs). These systems are also the target of high levels of antimicrobial use (AMU); however little is known about the profitability and sustainability of such systems. Since small scale farms are commercial enterprises, this knowledge is essential in order to develop successful strategies aimed at curbing excessive AMU. Using longitudinal data from 203 randomly selected small-scale (100-2,000 chickens) native chicken flocks raised in 102 farms in Dong Thap province (Mekong Delta, Vietnam), we investigated the financial and economic parameters of such systems and the main constraints to their sustainability. Feed accounted for the largest financial cost (flock median 49.5% [Inter-quartile range (IQR) 41.5-61.8%]) of total costs, followed by day-old-chicks (DOCs) (median 30.3% [IQR 23.2-38.4%]), non-antimicrobial health-supporting products (median 7.1% [IQR 4.7-10.5%]), vaccines (median 3.1% [IQR 2.2-4.8%]), equipment (median 1.9% [IQR 0.0-4.9%]) and antimicrobials (median 1.9% [IQR 0.7-3.6%]). Excluding labor costs, farmers achieved a positive return on investment (ROI) from 120 (59.1%) flocks, the remainder generated a loss (median ROI 124% [IQR 36-206%]). Higher ROI was associated with higher flock size and low mortality. There was no statistical association between use of medicated feed and flock mortality or chicken bodyweight. The median daily income per person dedicated to raising chickens was 202,100 VND, lower than alternative rural labour activities in the Mekong Delta. In a large proportion of farms (33.4%), farmers decided to stop raising chickens after completing one cycle. Farmers who dropped off chicken production purchased more expensive feed (in 1,000 VND per kg) (11.1 [10.6-11.5] vs. 10.8 [10.4-11.3] for farms that continued production (p=0.039) and experienced higher chicken mortality (28.5% [12.0-79.0%] vs. 16 [7.5-33.0%] (p=0.004). The turnover of farmers raising chickens in such systems represents a challenge in targeting messages on appropriate AMU and on chicken health. In order to ensure sustainability of small-scale commercial systems, advisory services need to be available as farmer initiate new flocks, and support them in the early stages to help overcome their limited experience and skills. This targeted approach would support profitability whilst reducing risk of emergence of AMR and other disease problems from these systems.

## INTRODUCTION

In Vietnam and many other low- and middle-income countries (LMICs) raising small-scale chicken flocks is a common activity that contributes to the income of many rural households. In addition to providing food security and income, such farming systems help promoting community relations (Alders and Pym, 2009). The Mekong Delta region of Vietnam (human population 21.5 millions in 2019) represents ∼13% of the total national chicken meat output (840,000 tons in 2018) (General Statistics Office Of Vietnam, 2018). Chicken production in the area is predominantly semi-intensive (including backyard and small-scale), and is typically based on slow-growing native breeds (Duc and Long, 2008; Lan Phuong et al., 2015). Production is however hampered by a high incidence of parasitic (Nguyen T. B. Van et al., 2020), viral and bacterial diseases (Nguyen Thi Bich Van et al., 2020), often resulting in high mortality losses (Carrique-Mas et al., 2019b). Furthermore, levels of AMU in these systems are known to be particularly high. A recent survey in the area reported that, on average, farmers administered 323.4 (SEM ±11.3) mg of antimicrobial active ingredients (AAI) per kg of chicken sold. In addition, chickens are often raised on commercial medicated feed (estimated to amount to ∼85 mg/kg chicken sold) (unpublished). The use of a total of 42 different antimicrobials, many of which are of critical importance by the World Health Organization (WHO) has been described in chicken flocks in the area (Cuong et al., 2019). Chicken farmers often use antimicrobials with the aim of preventing disease, especially during the brooding period, since antimicrobials are viewed as a cheaper alternative than other disease control measures (Truong et al., 2019). Antimicrobials are typically sold over the counter and prices are generally very low (estimated in ∼0.40 cents of 1 USD per daily dose administered to a 1kg chicken) (Dung et al., 2020).

Economic analyses of broiler production systems have been performed in Pakistan and Indonesia (Afzal and Khan, 2017; Coyne et al., 2020b). However, there are no published data quantifying the financial flows within small-scale chicken farms raising native chickens that are so common in Vietnam and other Southeast Asian countries. These farming units are typically smaller than their counterpart broiler farms, feed/water dispensation is manual, and birds are always raised at ambient temperatures. A main challenge for studying these systems is the lack of record-keeping practices in many units (Coyne et al., 2019). Using economic disease and production data from cohorts of small-scale chicken flocks raised in the Mekong Delta over 18 months followed up from day-old to slaughter age, we characterized the cost structure of such systems with the aim of quantifying the fraction spent on antimicrobials and other key inputs. The integration of data on feed medication and AMU allowed use to investigate the impact of these parameters on enterprise productivity. An understanding of the economic parameters that underpin small-scale production systems is a pre-requisite for developing and implementing strategies aiming at improving animal health, whilst reducing excessive AMU in Vietnam and elsewhere in Southeast Asia.

## MATERIALS AND METHODS

### Study area and data collection

The study was conducted in Cao Lanh and Thap Muoi districts, Dong Thap province (Mekong Delta, Vietnam) from October 2016 to March 2018. The human population of the province (2017) was 1.69 million. The population densities are 500 people/km^2^ and 127 chicken/km^2^ (General Statistics Office (GSO), 2018) (Sub-Department of Animal Health of Dong Thap, 2017). The study was based on data collected during the baseline phase of an intervention study (Carrique-Mas and Rushton, 2017). Randomly selected owners of chicken farms were drawn from the official census and were invited to enroll in the study. We aimed at recruiting farms raising flocks with >100 heads each, managed as all-in-all-out (i.e. single age). Enrolled farmers that consented to the study were provided with diaries and were requested to weekly record information on disease, mortality, source of day-old chicks (DOCs), types and amounts of feed used and health-related products (antimicrobials, vaccines, antiparasitic drugs, disinfectants), as well as any equipment purchased. The costs incurred by farmers, the weight of chickens at point of sale and the income generated from chicken sales were also recorded. Farms were visited by trained field researchers to review the information collected by the farmer on four different occasions over each production cycle. These data were transferred onto a questionnaire and were then uploaded to a central database using a web application for further analysis. This study was granted ethics approval by the Oxford Tropical Research Ethics Committee (OxtTREC) (Ref. 5121/16) and by the local authorities (People’s Committee of Dong Thap province).

### Data analyses

The sum of all financial costs incurred in procuring DOCs and raising the flocks until slaughter age were computed as input data; output data consisted of the revenues derived from the sale of chickens. For each flock we computed the difference between inputs and outputs, excluding the costs of labour. Labour costs were analyzed separately, since they were an opportunity cost, not a financial one. We calculated the return on investment value without labour costs (ROI) for each flock raised (Equation 1) (Zamfir et al., 2016). The ROI value included three range, < 0 (negative value and negative profit); <0 - ≤100 (positive value and negative profit – invest 1 VND but return less than or equal to 1 VND -); >100 (positive value and positive profit – invest 1 VND and return more than 1 VND-).

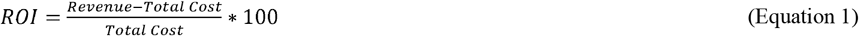

Financial costs and revenues were expressed ‘per chicken sold’ at the end of each production cycle.

Farm- and flock-related factors influencing the ROI excluding labour for each flock produced were investigated by building a linear mixed random-effects model. Farm-related variables included district location, owner’s gender, owner’s age, number of staff working in the farm (including the owner), experience in commercial poultry farming (in years) and education achievement of the owner. Flock-related variables were flock size (No. chickens purchased), duration of the production cycle (in weeks), number of sources of DOCs and their cost, feed type (commercial feed, locally-sourced) and price (per kg), percent of weeks consuming commercial medicated feed, average number of daily doses of antimicrobial administered to 1 kg of live chicken per 1,000 kg chicken-days (ADD_kg_ per 1,000 kg chicken-days) (Phu et al., 2020), number of antimicrobial-containing products used, number of vaccines (pathogens) per flock, flock cumulative mortality over the production cycle (as percent of chickens purchased), cumulative mortality from week 9 (as percent of chickens purchased), flocks in farms raising >1 flock simultaneously, flocks in farms also raising non-chicken species, flocks where farmer purchased new equipment. All variables were tested as fixed effects, with ‘farm’ identity included as a random effect. Factors that were significant (p<0.20) in univariable analysis were included in multivariable analysis.

We investigated the potential association between all modelled variables (including weight of chickens at sale). Of particular interest was the association between (1) price of DOCs and (i) weekly mortality, (ii) duration of the production cycle and (iii) chicken weight at sale; (2) No. of ADD_kg_ (per 1,000 kg chicken-days) and (i) weekly mortality or (ii) flock size and (iii) chicken weight at sale; (3) percent of weeks on medicated feed and (i) weekly mortality, (ii) flock size and (iii) chicken weight at sale; and (4) weekly mortality and flock size. We related the income generated from the flocks to the total time spent by the farm owner and other staff (including relatives) tendering the flocks (Equation 2).

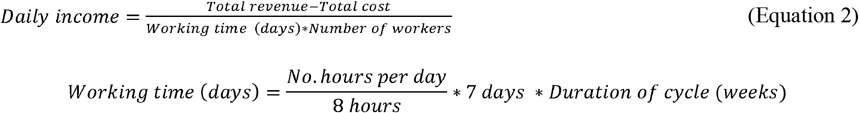

All financial revenues and costs were expressed in thousands (1,000s) of Vietnam Dong (VND). We investigated the differences between flocks that were and were not followed by a subsequent one within 8 months (i.e. continued/discontinued production) with respect to variables included in the previous analyses. The chosen criterion ensured that seasonal farmers (i.e. those that regularly raise only one cycle per year, typically for the annual Tet holiday) were not considered discontinued production. Pearson’s Chi-squared tests were used for proportions and (non-parametric) Wilcoxon rank sum tests were used for continuous data.

## RESULTS

### Description of study flocks

A total of 102 farms and 203 flock production cycles were investigated (Table 1). The median flock size was 300 [Inter-quartile range (IQR) 201-502]. A median of 2 [IQR 1-2] flocks were investigated per farm. Chicken flocks were raised over a median of 18 weeks [IQR 16-20]. The median cumulative mortality of flocks over the production cycle was 18% [IQR 8-40%]. Cumulative mortality reached 100% in 8 (3.9%) flocks that were affected by an outbreak of severe disease. The main financial revenue in these flocks derived from the sale of live chickens for meat. In all cases farmers collected manure (used litter) and feathers and were used to fertilize crops with no associated income.

**Table 1.**
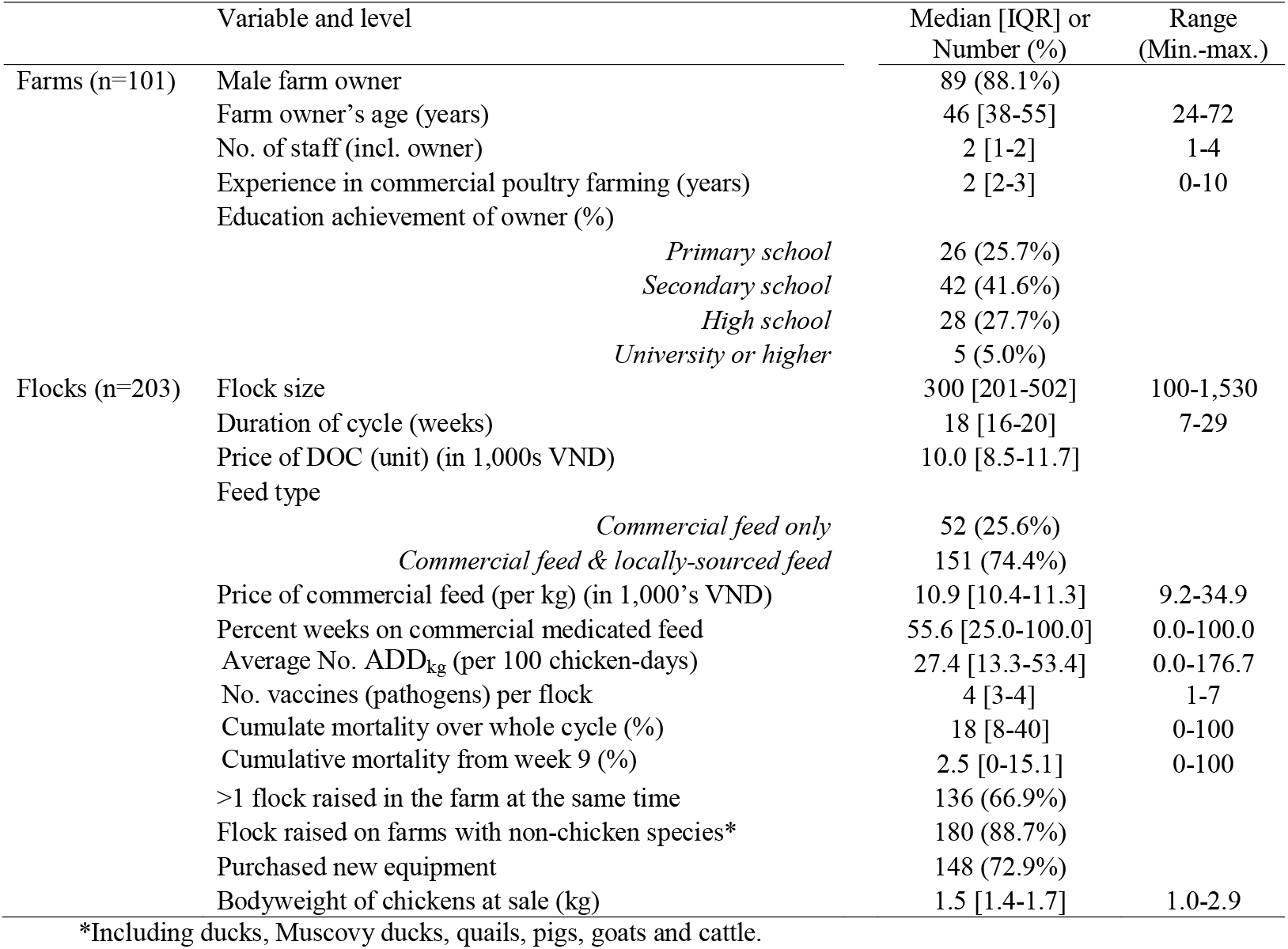
Description of key variables related to small-scale commercial chicken flocks raised in Dong Thap province (Mekong Delta, Vietnam).

### Cost structure over the flock production cycle

The median financial cost across flocks (in 1,000s VND) incurred in raising one chicken from day-old to slaughter was 48.3 [IQR 36.8-78.0]. Feed cost (commercial and locally-source feed) accounted for a median of 49.5% [IQR 41.5-61.8] of the total cost, followed by DOCs (median 30.3% [IQR 23.2-38.4]), non-antimicrobial health-supporting products (vitamins, anti-parasitic drugs) (median 7.1% [IQR 4.7-10.5]), vaccines (median 3.1% [IQR 2.2-4.8]) and other costs (median 1.9% [IQR 0.0-4.9]) (including equipment, litter, electricity and disinfectants). Expense on antimicrobials accounted for a median of 1.9% [IQR 0.7-3.6] of total costs. The median revenue obtained per chicken sold was 108.7 thousand VND [IQR 99.4-121.7]. Flocks were raised on commercial feed containing antimicrobials over a median of 55.6% weeks [IQR 25.0 – 100.0]. The mean price (by product) of commercial medicated feed was estimated in 11.2 (Standard Deviation (SD) ±2.5 1,000s VND/kg, higher than the price of non-commercial feed (11.1 SD ±0.4 per kg) (Welch’s t test 2.19, p=0.028). The median ROI across all flocks was 124% [IQR 36-206%]; therefore, for each VND invested there was a return of VND 1.24. For 120/203 (59.1%) flocks, farmers obtained a positive profit (ROI>100%), whereas for 83 (40.9%) farmers obtained negative profit (ROI<100%) (i.e. financial losses). The main cost categories sorted by flock ROI by flock are presented in Figure 1. Generally, higher ROI was associated with a smaller fraction of feed costs.

**Figure 1.**
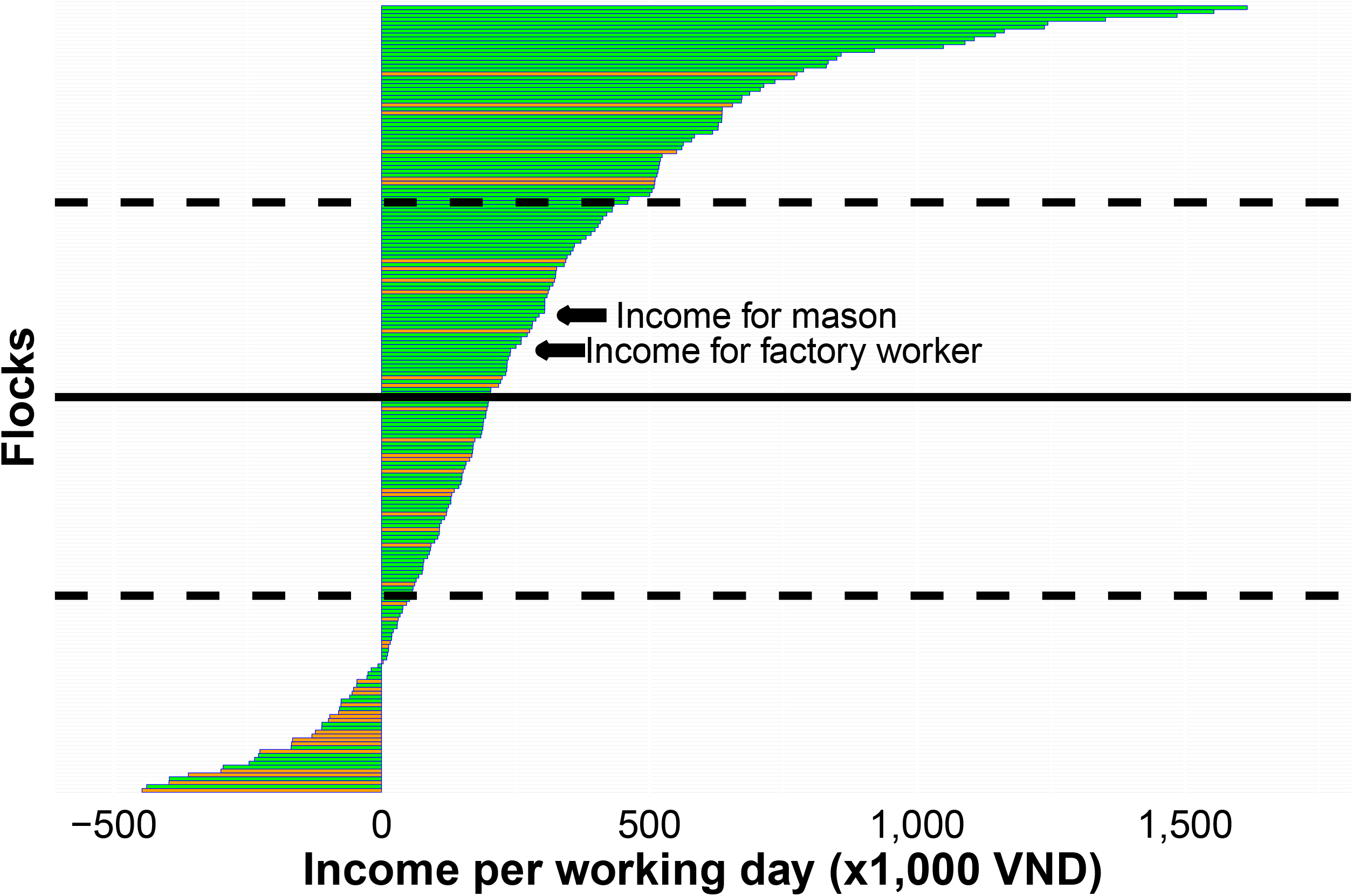
Graphic representation of the cost structure of small-scale chicken flocks raised in Dong Thap province, sorted by ROI, and stratified by four categories: (1) Feed; (2) Day-old chicks; (3) Antimicrobials and vaccines; (4) Other costs. Solid line (mean); dashed line (median). Flocks that make a positive profit have ROI>0.

The correlation between all modelled variables (including weight of chickens at sale, not modelled) is displayed in Figure 2. There was no association between the cost of DOCs (per unit purchased) and cumulative mortality (Pearson’s correlation coefficient *r*=-0.11, p=0.093) or the duration of the flock cycle (*r*=-0.01; p=0.877). There was a negative correlation between the cost of DOCs and chicken weight at sale (*r*=-0.14, p=0.041), as well as between the total number of ADD_kg_ administered and flock size (*r*=-0.19, p=0.020).

**Figure 2.**
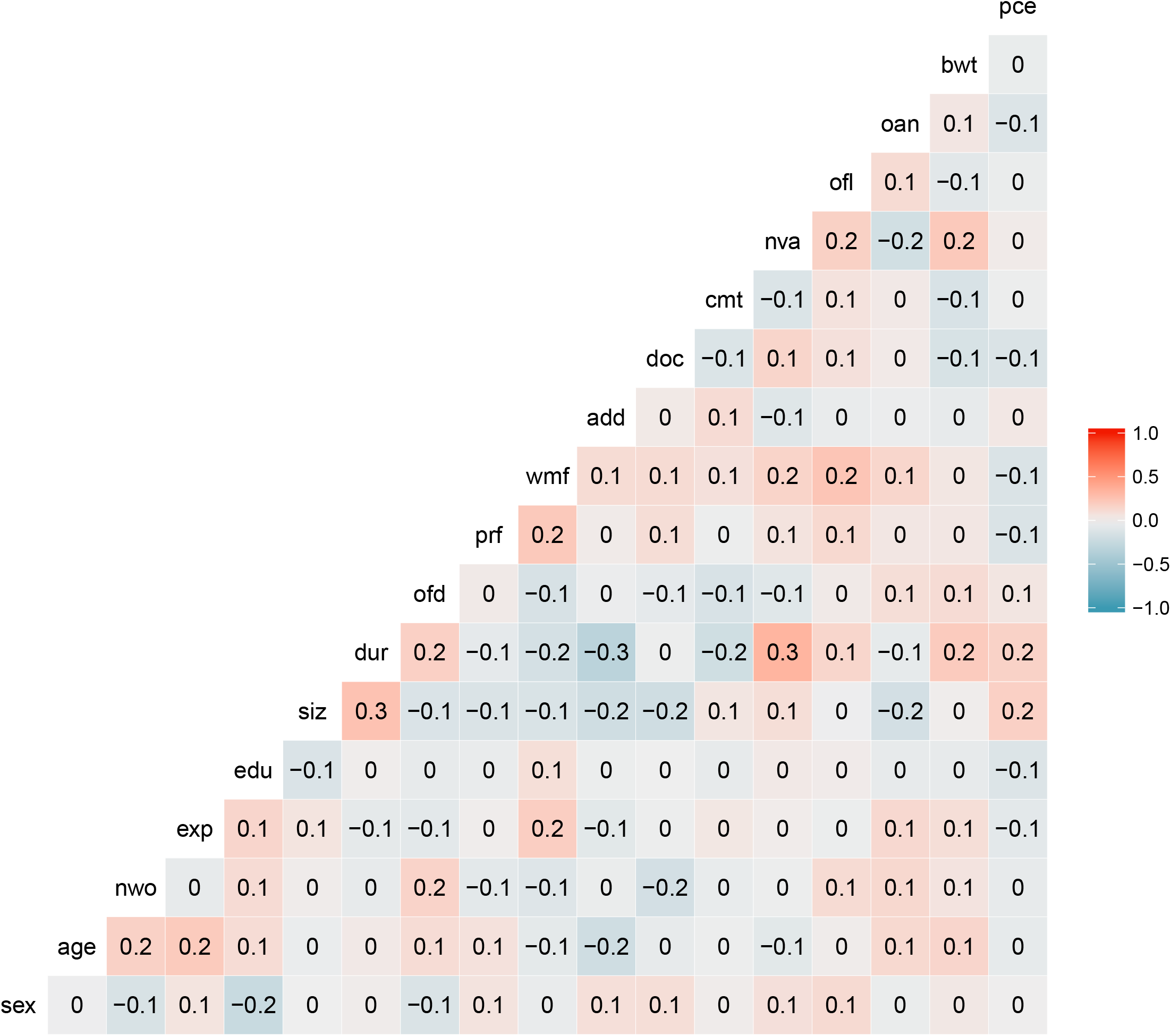
Correlation matrix of study variables. *sex*: owner’s gender, *age:* owner’s age; *nwo:* number of workers; *exp:* experience in poultry farming (years); *siz:* flock size; *dur:* length of cycle; *ofd:* usage locally-sourced feed; *prf:* feed price; *wmf:* percent weeks on medicated feed; *add:* average weekly antimicrobial daily dose; *doc:* price DOCs; *cmt:* flock cycle cumulative mortality; *nva:* number of pathogens vaccinated; *ofl:* other flock raised in farm; *oan:* other animal raised in farm; *bwt:* average weight of chicken; *pce:* purchase of new equipment.

### Factors associated with ROI excluding labour costs

The ROI values were square root-transformed in order to improve their distribution’s normality for subsequent modelling. Only two factors were independently associated with ROI (inverse association) were flock size (*β* =0.08, p<0.001), and cumulative mortality (*β* =-1.45, p<0.001) (Table 3). As expected, the average weight of chickens sold was strongly associated with flock ROI (not included in the multivariable model) (*β* =0.51, p=0.001). The predicted outcomes for different values of the two significant variables are given in Supplementary Material 1. Mortality was the single most influential driver of ROI.

**Table 2.**
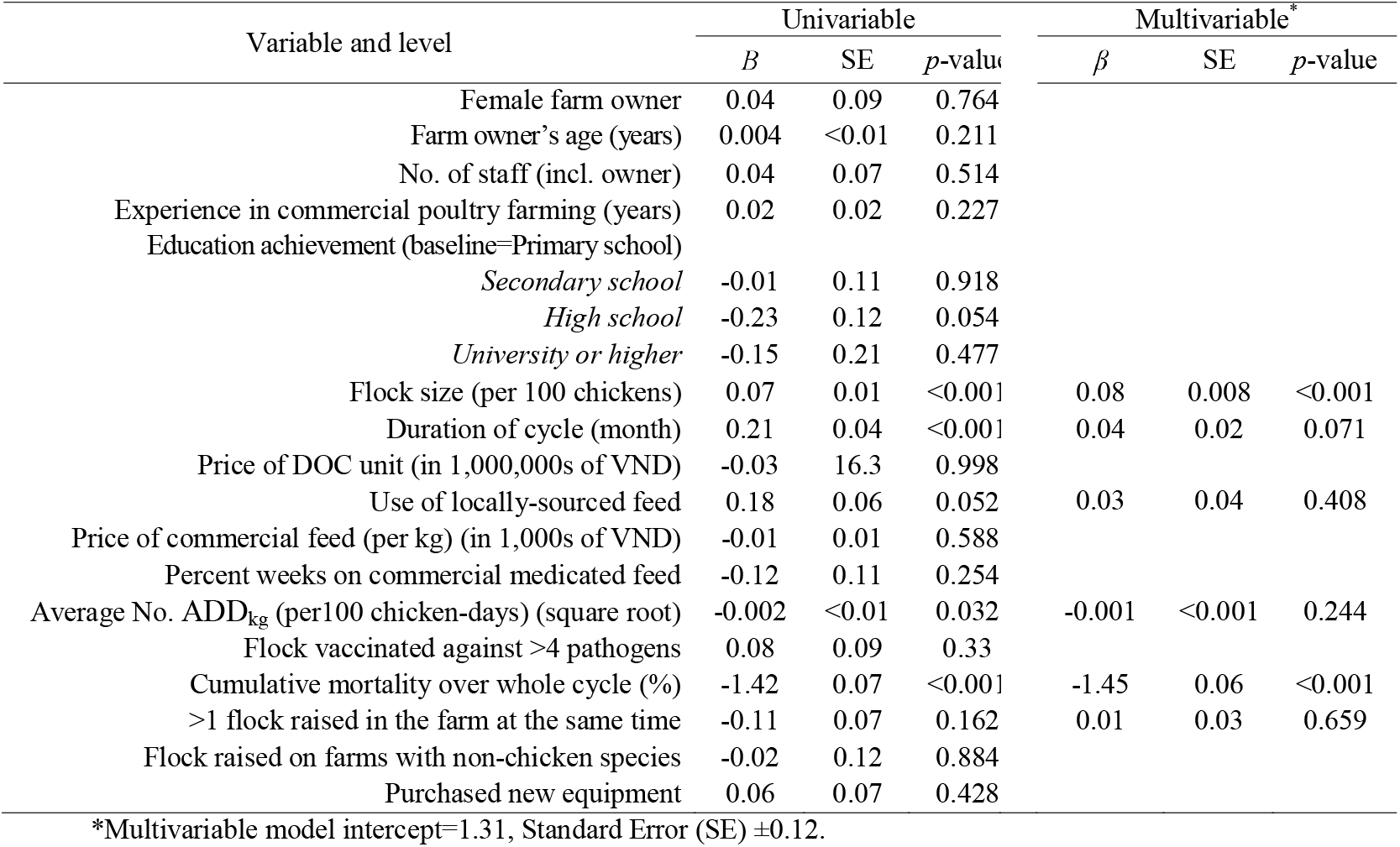
Linear mixed models investigating factors associated with ROI of raising small-scale meat chicken flocks.

| Variable and level | Univariable |  |  | Multivariable* |  |  |
| --- | --- | --- | --- | --- | --- | --- |
| | *B* | SE | *p*-value | $\beta$ | SE | *p*-value |
| Female farm owner | 0.04 | 0.09 | 0.764 |  |  |  |
| Farm owner's age (years) | 0.004 | <0.01 | 0.211 |  |  |  |
| No. of staff (incl. owner) | 0.04 | 0.07 | 0.514 |  |  |  |
| Experience in commercial poultry farming (years) | 0.02 | 0.02 | 0.227 |  |  |  |
| Education achievement (baseline=Primary school) |  |  |  |  |  |  |
| *Secondary school* | -0.01 | 0.11 | 0.918 |  |  |  |
| *High school* | -0.23 | 0.12 | 0.054 |  |  |  |
| *University or higher* | -0.15 | 0.21 | 0.477 |  |  |  |
| Flock size (per 100 chickens) | 0.07 | 0.01 | <0.001 | 0.08 | 0.008 | <0.001 |
| Duration of cycle (month) | 0.21 | 0.04 | <0.001 | 0.04 | 0.02 | 0.071 |
| Price of DOC unit (in 1,000,000s of VND) | -0.03 | 16.3 | 0.998 |  |  |  |
| Use of locally-sourced feed | 0.18 | 0.06 | 0.052 | 0.03 | 0.04 | 0.408 |
| Price of commercial feed (per kg) (in 1,000s of VND) | -0.01 | 0.01 | 0.588 |  |  |  |
| Percent weeks on commercial medicated feed | -0.12 | 0.11 | 0.254 |  |  |  |
| Average No. ADD<sub>kg</sub> (per100 chicken-days) (square root) | -0.002 | <0.01 | 0.032 | -0.001 | <0.001 | 0.244 |
| Flock vaccinated against >4 pathogens | 0.08 | 0.09 | 0.33 |  |  |  |
| Cumulative mortality over whole cycle (%) | -1.42 | 0.07 | <0.001 | -1.45 | 0.06 | <0.001 |
| >1 flock raised in the farm at the same time | -0.11 | 0.07 | 0.162 | 0.01 | 0.03 | 0.659 |
| Flock raised on farms with non-chicken species | -0.02 | 0.12 | 0.884 |  |  |  |
| Purchased new equipment | 0.06 | 0.07 | 0.428 |  |  |  |

\*Multivariable model intercept=1.31, Standard Error (SE) ± 0.12.

**Table 3.**
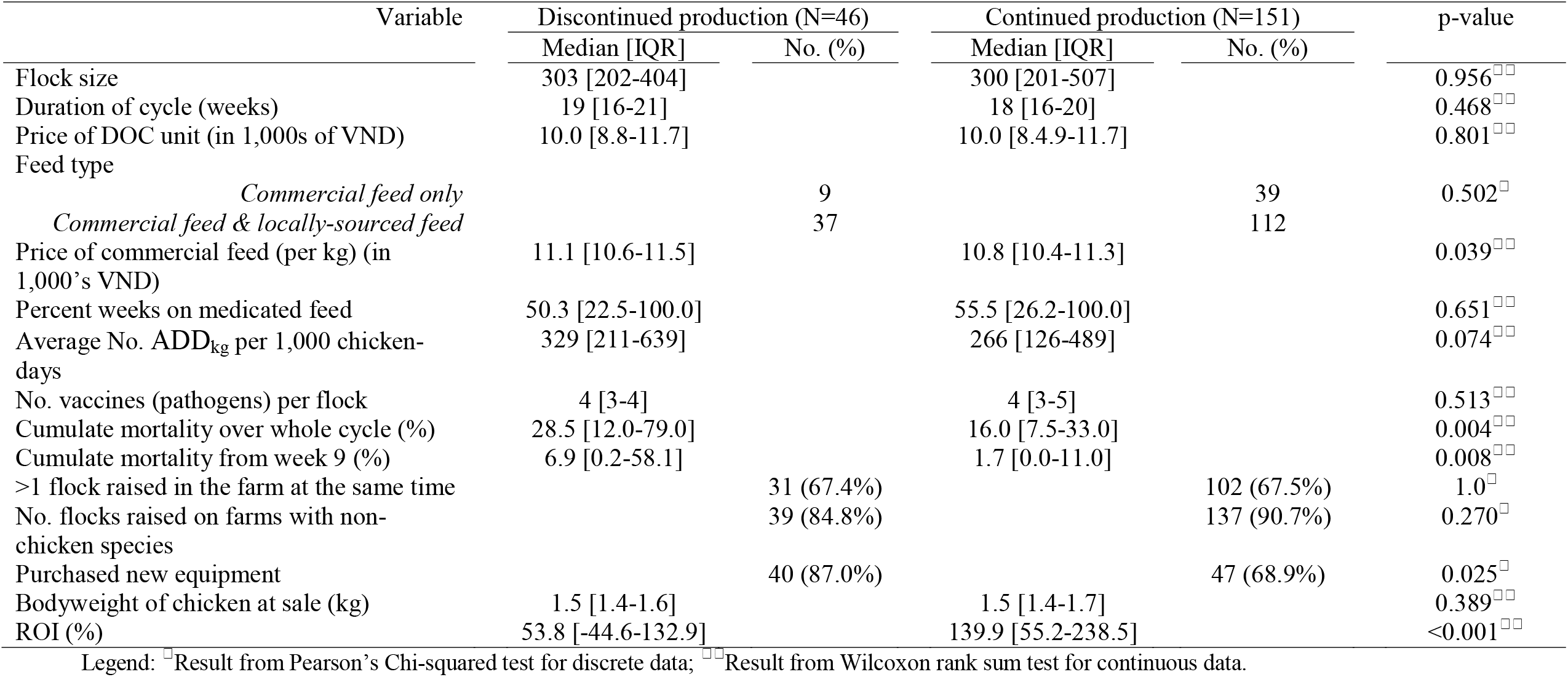
Description of farm/flocks that discontinued/continued chicken production after one flock.

### Income generated per day of labour

A median of 399 [IQR 266-613] person-hours (equivalent to 49.8 [IQR 33.3-76.5] working days were employed to raise one flock, and a median of 2.3 [IQR 1.3-4.4] person-hours per chicken raised. The median daily income per person in chicken production was 202.1 thousands of VND [IQR 56.5-461.0]. However, for 33 (16.2%) flocks farmers labour income was negative. There was a high positive correlation between income per working day and flock size (*r*=0.213, p<0.01) (Figure 3).

**Figure 3.**
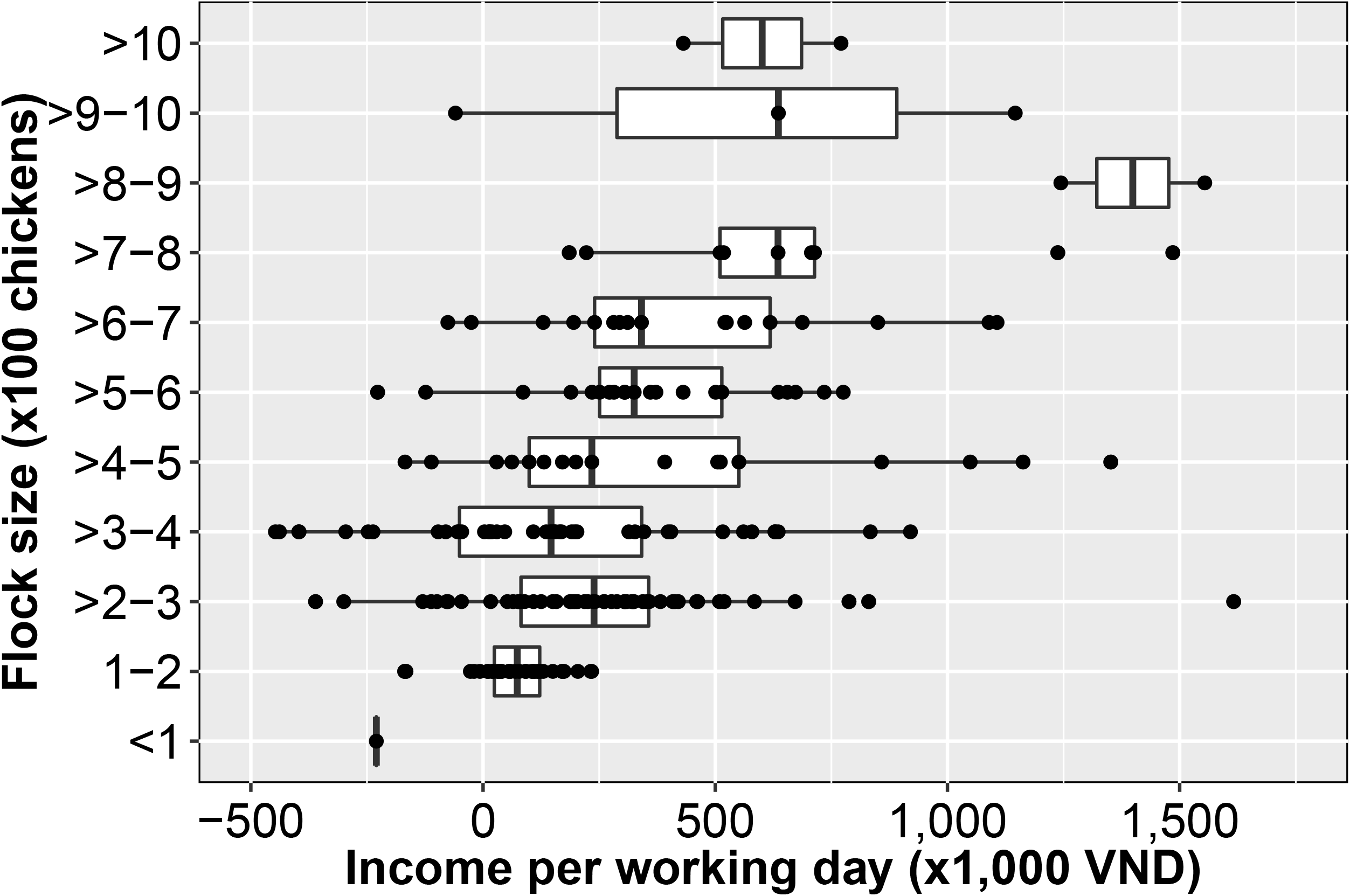
Income generated per day of labour according to flock size.

**Figure 4.**
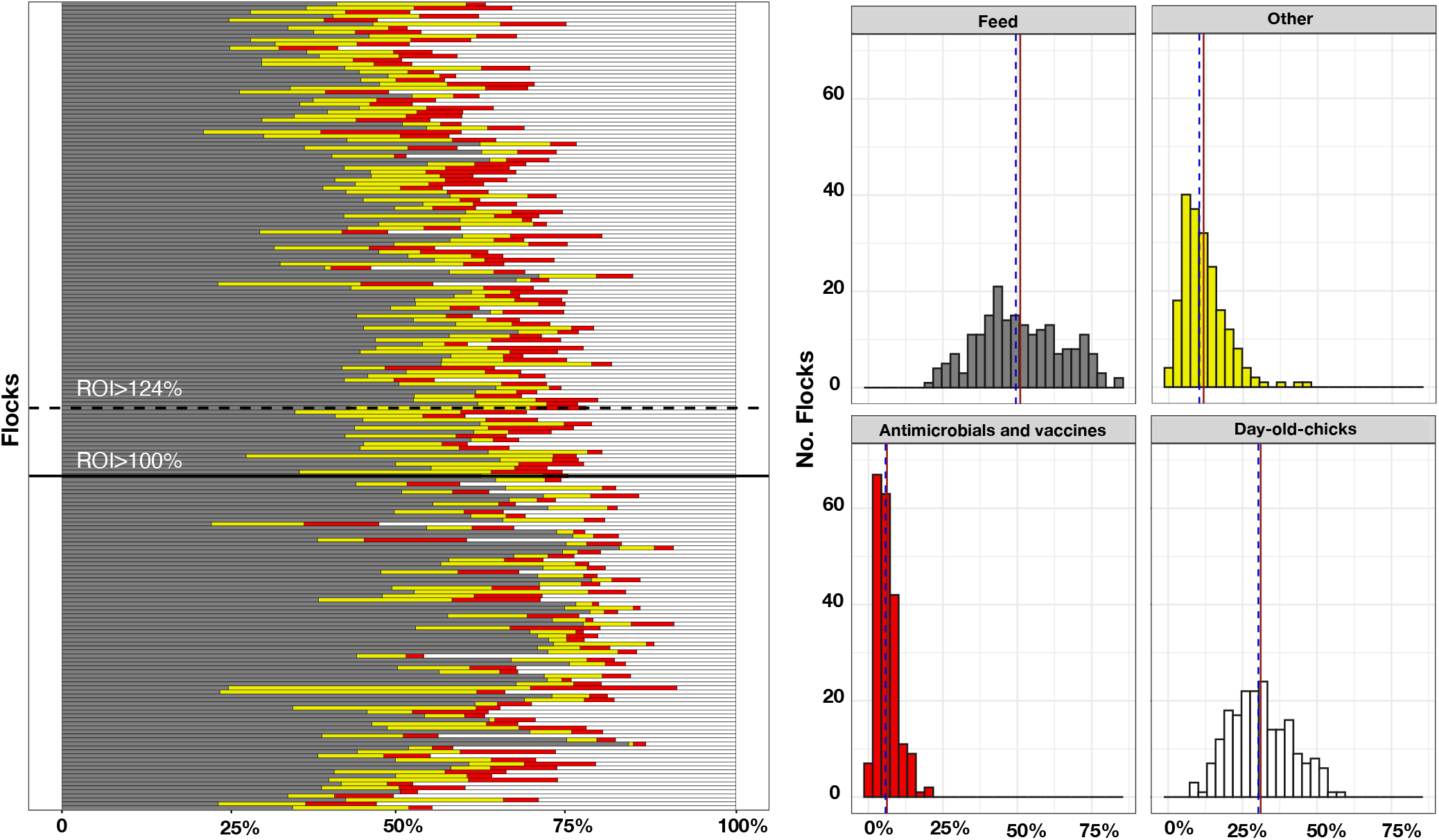
Distribution of income generated per day of labour among flocks. Solid line (mean); dashed lines (1^st^ and 3^rd^ quartile); green (farmers that continued production); orange (farmers that discontinued production)

### Farms that discontinued chicken production

Of 197/203 (97%) flocks that could be evaluated for the criterion of whether or not farmers continued raising chickens within 8 months after their sale, 46 (33.4%) were not followed up by a subsequent flock (i. e. discontinued farming). Table 3 shows the differences with regards to all variables investigated. Farmers that did not raise further flocks had purchased feed prices (per kg) at a higher cost compared with flocks in farms that continued production (11.1 [10.6-11.5] 1,000 VND vs. 10.8 [10.4-11.3]; p=0.039). In addition, these farmers that discontinued production also experienced higher mortality in their flocks (28.5% [12.0-79.0%] vs. 16% [7.5-33.0%]; p=0.004). As expected, the ROI of flocks for flocks followed up were also higher (53.8% [-44.6-132.9%] vs. 139.9% [55.2-238.5%]) (Table 3).

## DISCUSSION

Our study shows that, in the Mekong Delta of Vietnam, raising flocks of 100-2,000 meat chickens is generally profitable (median 1.24 VND returned per VND invested). The financial profit generated from these systems (per chicken produced) generally increased with flock size. However, in 40.9% of cases farmers incurred in financial losses. The main explanatory factors for the losses identified were a small flock size and high mortality. The (high) observed cumulative mortality (median 18%) is likely to reflect a high incidence of bacterial, viral and parasitic diseases identified in the area (Carrique-Mas et al., 2019b; Nguyen T. B. Van et al., 2020; Nguyen Thi Bich Van et al., 2020). This cumulative mortality reached 28% in flocks in farms that discontinued chicken production.

DOCs in our study were generally more expensive (median 30.3% of input costs) compared with Indonesian (25% costs) and Pakistani (22-29%) broiler flocks (Afzal and Khan, 2017; Coyne et al., 2020b). This higher price of DOCs in our flocks is probably due to the absence of commercial hatcheries in the study province (Dong Thap), with DOCs typically delivered through a complex network of intermediate traders. We did not attempt to characterize the breed identity of study flocks due to its complexity. The choice of adequate breed lines should however be important in order to maximize production in these small-scale systems.

Our study indicates that commercial feed was still the greatest single most important financial expense incurred to raise chicken flocks (median 49.5% of all costs). In Pakistani broiler flocks feed represented 58.1-63.6% depending on farm type (Afzal and Khan, 2017), and in Indonesia ∼70% (Coyne et al., 2020b). It is possible that some of savings in our study flocks were due to the provision of locally sourced feed (local plants, rice gain), normally at no cost. Disinfectants account for a very small fraction since in the area as they are often provided by the veterinary authorities (Dong Thap Sub-Department of Animal Health) free of charge. Our results indicate that, overall, the cost (per kg) of medicated feed was marginally higher (∼1%) than that of non-medicated feed; however, we could not demonstrate any impact of medicated feed on bodyweight and/mortality. A total of eight antimicrobial active ingredients (AAIs) were identified from the examination of the labels of these commercial feed formulations (avilamycin, bacitracin, chlortetracycline, colistin, enramycin, flavomycin, oxytetracycline, virginamycin). However only three AAIs, chlortetracycline, bacitracin and enramycin, amounted to >90% of total use (data not shown). The evidence of the impact of antimicrobials in feed (generally as antimicrobial growth promoters, AGP) on animal productivity has been mixed and highly variable depending on the studies and production types (Laxminarayan et al., 2015). In our study, a lower price of commercial feed was associated with continued chicken farming; this confirms that the affordability of this input is crucial to the sustainability of the production system.

We estimated that medicines accounted for ∼9% of total input costs, compared with 5-6% in Pakistan and ∼1.3% in Indonesia. However, in our study the fraction of antimicrobials to total medicine cost was relative small. In contrast, expense on vaccines was higher in our study flocks (3.1%) than in the Indonesian (0.8%) and Pakistani broiler studies (∼1.5) (Afzal and Khan, 2017; Coyne et al., 2020b). Vaccination of flocks against pathogens is a widespread practice in the Mekong Delta, with >85% flocks vaccinated against three or more pathogens, the most common being, in decreasing order, were Newcastle Disease, Highly Pathogenic Avian Influenza, Infectious Bursal Disease, fowl pox, fowl cholera and Infectious Bronchitis (data not shown).

Our results reflect a relatively small impact of AMU on overall financial costs (<2%) in spite of the high volume of antimicrobials used (previously described in detail using a number of metrics (Cuong et al., 2019). This is reflective of the generally low price of antimicrobials available to Mekong Delta farmers. For example, a daily dose of an average antimicrobial-containing product (per LJkg of chicken) has been estimated to retail at ∼0.40 cents of 1 US$, depending on product (Carrique-Mas et al., 2019a; Dung et al., 2020). Our findings are not dissimilar to costs of antimicrobials intended for small-scale pig farms in Vietnam (<2% total costs) (Coyne et al., 2020a).

In Vietnam, as well as in other ASEAN countries, livestock production systems have become more intensified in recent years in response to increased demand for animal protein (Jabbar, 2015; Soedjana and Priyanti, 2017). Small and medium-scale production systems such as those described in this study have accordingly become more prevalent and have generally become larger to improve competitiveness. The systems described here represent a transition from backyard to industrial production. However, an increase in flock presents associated challenges, since increased flock size normally entails a high risk of disease and mortality. A recent study has shown that levels of mortality are generally larger in larger flocks, as well as AMU (in terms of frequency) (Carrique-Mas et al., 2019b). In contrast with the findings of that study, we found that larger flock generally received fewer doses compared with small flocks. This is due to small flocks being more prone to overdosing due to incorrect preparation of the antimicrobial product (authors’ observation).

Despite the increased popularity of industrial broilers, meat from native breeds is still highly valued by Vietnamese consumers thanks to its distinct taste. Native chickens purchased at the farm gate (median price ∼70,000 VND per kg in our study) already reached a much higher retail price than broiler chicken meat purchased at retail (i.e. supermarket) (∼20,000-25,000 VND per kg) (Anonym, 2019). In Vietnam native chicken production still represents a majority of national chicken meat output: of 990,397 tonnes of chicken meat produced in 2019, 43.3% corresponded to industrial chicken production, the rest corresponding to meat from backyard and small scale production systems (Anonym, 2019).

The estimated daily income from raising chickens in our study (VND 202,120) was considerably lower than that generated from comparable occupations in the area, such as mason (VND ∼300,000 per day), or factory worker (VND ∼250,000 per day) (authors’ observation). A major challenge is the limited available land in this densely populated area and the competing income-generating activities (retail, rice, fruit, duck, pig and fish farming). Most households normally make up their income from a range of activities. Often they resort to other members in the family for support in raising chickens. For the individual farmer, raising livestock and poultry are often unstable activities due to the seasonality of production, market fluctuations and the regular incursion of infectious diseases. The recent 2019 incursion of African Swine Fever epidemic in Vietnam (Le et al., 2019) resulted in many pig farmers switching to chicken production, with an associated decline in prices of finished chicken. The necessities to undertake multiple occupations are not generally conducive to farmers becoming proficient in chicken husbandry.

## Conclusion

We confirmed considerable instability in small-scale chicken farming systems in the Mekong Delta, with many farmers ceasing production after one or two cycles. A higher profitability was attained with larger flock sizes and low mortality. A higher cost of DOCs was associated to increased mortality. In spite of a high frequency of AMU, expenses on antimicrobials accounted for <2% of total costs. We did not find any impact of medicated feed on overall mortality or productivity in flocks. In order to remain profitable, farm owners need to implement effective disease control practices. Establishing advisory services for farmers to help them improve their knowledge base on flock management and health issues, whilst reducing levels of AMU should be a priority for authorities and policy makers. This targeted approach would support profitability whilst reducing risk of emergence of AMR and other disease problems from small-scale production systems.

## Acknowledgements

This work was funded by the Wellcome Trust through an Intermediate Clinical Fellowship awarded to Juan J. Carrique-Mas (Grant Reference Number 110085/Z/15/Z). We are grateful to the Sub-Department of Animal Health of Dong Thap and Nong Lam University for supporting field work.

