## Supplementary 1 for "Small-scale commercial chicken production: A risky business for farmers in the Mekong Delta of Vietnam"

**Supplementary Material 1**

Predictions for ROI derived from the final multivariable model for three levels of each explanatory variable (25% quartile, median and 75% quartile). Calculations for each variable were done after fixing the remaining variable at its median value. Model equation: ROI = 1.5 + 0.09*flock size (per 100 chickens) -1.48*Cumulative mortality (percent).

|  | 25% quartile | Median | 75% quartile |
| --- | --- | --- | --- |
| Flock size (201, 300, 502) | 114.7% | 125.0% | 148.8% |
| Mortality (8%, 18%, 40%) | 143.5% | 123.7% | 90.9% |
